# Using Repeated Lysis Steps Fractionates Between Heterotrophic and Cyanobacterial DNA Extracted from Xenic Cyanobacterial Cultures

**DOI:** 10.1101/2024.08.22.609136

**Authors:** Alexis D. Wagner, Mohammed M. A. Ahmed, Victoria Starks, Paul D. Boudreau

## Abstract

Extracting DNA from cyanobacteria can be a challenge because of their diverse morphologies, challenging cellular structure, and the heterotrophic microbiome often present within cyanobacterial cultures. As such, even when our DNA yields are sufficient for sequencing, the percentage of reads coming from the cyanobacterial host can be low, leading to incomplete genomes spread across several scaffolds. In this research, we optimized a DNA isolation protocol using three iterative cell lysis steps to enrich the portion of DNA isolated coming from the cyanobacterial host rather than the heterotrophic microbiome. In order to utilize in-house nanopore sequencing, we faced a challenge in that our lysis protocol led to DNA shearing and a lower molecular weight DNA extract than is suitable for this sequencing technology. As such we used two bead-based size selection steps to remove shorter molecules of DNA before nanopore sequencing. EPI2ME analysis of the processed reads from the iterative lysis steps showed that in the first lysis the heterotrophic microbiome could make up more than half of all reads, but with each lysis the proportion of reads coming from these other species decreased. Using our iterative lysis protocol, we were able to sequence the genomes of two cyanobacteria isolated from fresh water sources around northern Mississippi, namely *Leptolyngbya* sp. BL-A-14 and *Limnothrix* sp. BL-A-16. The genomes of these isolates were assembled as closed chromosomes of 7.2 and 4.5 Mb for *Leptolyngbya* sp. BL-A-14 and *Limnothrix* sp. BL-A-16, respectively. Because some cyanobacteria have symbioses with their heterotrophic microbiome it is not always possible to prepare axenic cultures of these organisms, we hope our approach will be useful for sequencing xenic cultures of cyanobacteria, but we can also imagine applications in studying this microbiome specifically by focusing sequencing efforts on the first fraction.

## Introduction

Found in diverse environments around the world, cyanobacteria are well-known for their production of a vast array of secondary metabolites including many leads in drug discovery efforts.[1–3] Due to the diversity of these chemical scaffolds, cyanobacteria are of interest in the biotechnological, industrial, agricultural, and pharmacological fields.[4] Although they have many beneficial implications, they are also known producers of harmful algal blooms that can lead to ecosystem destruction, wildlife death, and human health impacts.[5–8] In order to better understand the metabolic capabilities and natural product potential of these organisms, there is a great deal of interest in whole genome sequencing cyanobacteria.[3,9] These genomic datasets can be mined for biosynthetic gene clusters responsible for natural product production because these clusters are often highly organized within a genome and, even for diverse chemical scaffolds, often share common core biosynthetic genes.[10,11] However, cyanobacteria have multiple features which make them challenging substrates for DNA isolation: They grow with different morphologies, some are single cellular some are quasi-multicellular filaments that form macroscopic assemblages;[12,13] cyanobacteria possess specialized plant-like cell walls that cause difficulty for traditional chemical lysis protocols and produce enzyme inhibiting secondary metabolites which can contaminate the DNA, these are both challenges for some common protocols and commercialized bacterial DNA isolation kits.[12,14–17] Their large cells (large relative to most bacteria) also serve as a scaffold for a microbiome community of heterotrophic bacteria, the DNA of which will also be collected in most workflows.[14,18–20]

There are DNA extraction protocols that have been developed for cyanobacteria specifically, but they are often targeted for one specific morphology or species.[21] These protocols come with their own challenges, approaches using phenol/chloroform-based DNA extractions are popular;[14–17,21–23] but this mixture is highly toxic, making this method less desirable (especially for teaching labs with undergraduates), and much research effort has been expended to avoid these reagents.[24–28] In this research, we worked to develop a DNA extraction method that was applicable to multiple cyanobacterial morphologies even in the presence of a complex microbiome of heterotrophic bacteria. To do so we adapted the Zymo E.Z.N.A.® Plant DNA Kit, using repeated, increasingly severe, lysis steps. In sequencing these different DNA fractions, we show that the cyanobacterial cells survive (or at least their DNA is not easily recovered in) the initial lysis step, but many heterotrophs do not; allowing the proportion of cyanobacterial reads to increase in the later lysis steps so that better cyanobacterial genomes can be assembled from this data.

## Results and Discussion

### Isolation of cyanobacteria from fresh water sources across northern Mississippi

We collected water samples from water bodies in Northern Mississippi, placed them on ice, and brought back to the laboratory for processing. BL-A-14 was isolated from a water sample taken from Enid lake, while BL-A-16 was isolated from a water sample collected from the Tennessee-Tombigbee waterway. Upon returning to the laboratory the water was plated directly on freshwater BG-11 (FW-BG-11) agar plates,[29] then left under grow lights until cyanobacterial colonies were observed. These colonies were picked onto fresh plates until a single morphology of photoautotroph was observed, from this plate a colony was picked into 50 mL liquid freshwater BG-11 media. The liquid cultures have been grown in continuous liquid culture in our laboratory ever since.

### Initial efforts via Illumina sequencing with a commercial vendor

We prepared DNA extractions from our xenic cultures of cyanobacteria using the commercial E.Z.N.A. Plant DNA kit from OMEGA Bio-Tek. When submitted to Genewiz (a commercial vendor) for Illumina sequencing we got high quality read data back, but the resulting genomic assemblies contained hundreds of contigs (669 for BL-A-14 and 263 for BL-A-16) with N_50_’s in the tens of thousands of base pairs (21,192 for BL-A-14 and 86,514 for BL-A-16). Based on GC content and BLAST comparison of open reading frames within these sequences we suspected they came from both the cyanobacterial target and its heterotrophic microbiome. A result which highlights the challenge of working with xenic cultures, where even a DNA isolate that appears to have good purity, yield, and concentration can be confounded by the presence of DNA from heterotrophic bacteria.

### Iterative lysis steps to isolate DNA with reduced heterotrophic DNA contamination

We subsequently prepared three separate DNA samples from our cultures using the same Plant DNA Kit but modifying the lysis step to afford a brief 10 min lysate, then collecting the residual cell pellet to isolate another lysate after 20 min of lysis, and finally collecting a lysate on the residual cell pellet from the previous step once again with a one-hour lysis (Figure 1). These three lysates were then processed to isolate three separate DNA preparations. Before a sequencing run using the Oxford’s nanopore technology, the DNA had to be size-selected as our lysis product generally lead to significant sheering of the DNA; using two magnetic bead-based size selection washes with SERAMAG beads we were able to enrich for the longer reads sufficient to run on a nanopore cell. We then submitted basecalled reads from sequencing the three lysis fractions of BL-A-16 on nanopore flongle flow cells to analysis via Oxford’s EPI2ME’s “What’s In My Pot” (WIMP) tool. For the three BL-A-16 fractions A, B, and C, the data showed 54%, 17%, and 9% respectively of all reads coming from a single heterotrophic genus, *Pseudomonas* (Figure 2). The WIMP tool was less accurate at assigning reads from the cyanobacterium, some of these reads were even called as *Homo sapiens*, when no human data was ever seen in our final assemblies (as expected!). However, we saw fewer heterotrophic reads after each lysis step supporting our hypothesis that our early mild lysis steps are better at popping open heterotrophic cells while the cyanobacteria cells remain intact in the pellet until the later lysis steps.

**Figure 1.**
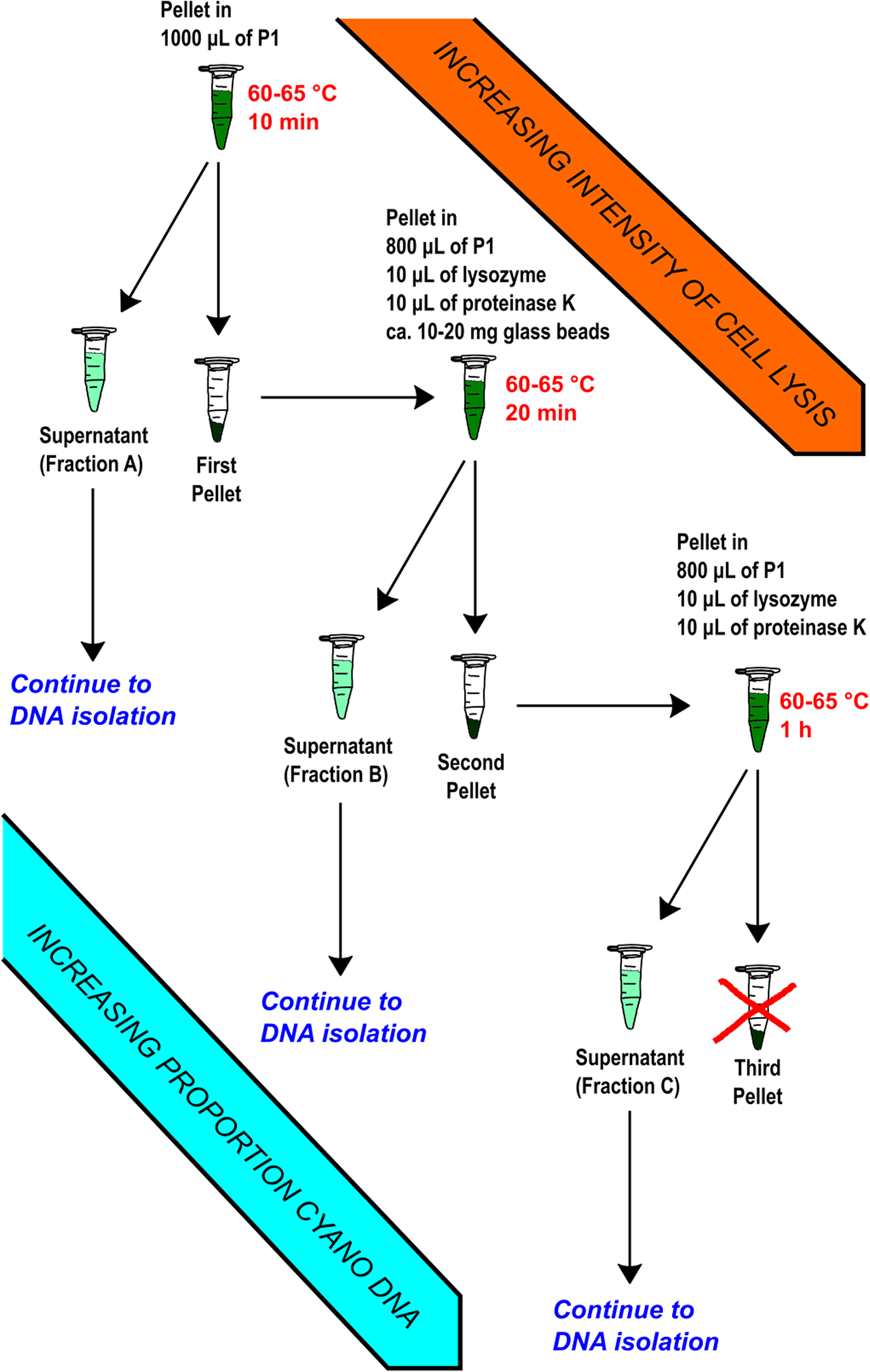
The iterative lysis workflow. The initial lysis step is incomplete, after collecting the pellet there is still DNA available to isolate in the solid cell matter. By repeating the lysis steps more vigorously, more DNA can be collected. In addition, because the cyanobacteria are generally more difficult to lyse than heterotrophic bacteria, the proportion of cyanobacterial DNA increases in each lysis fraction.

**Figure 2.**
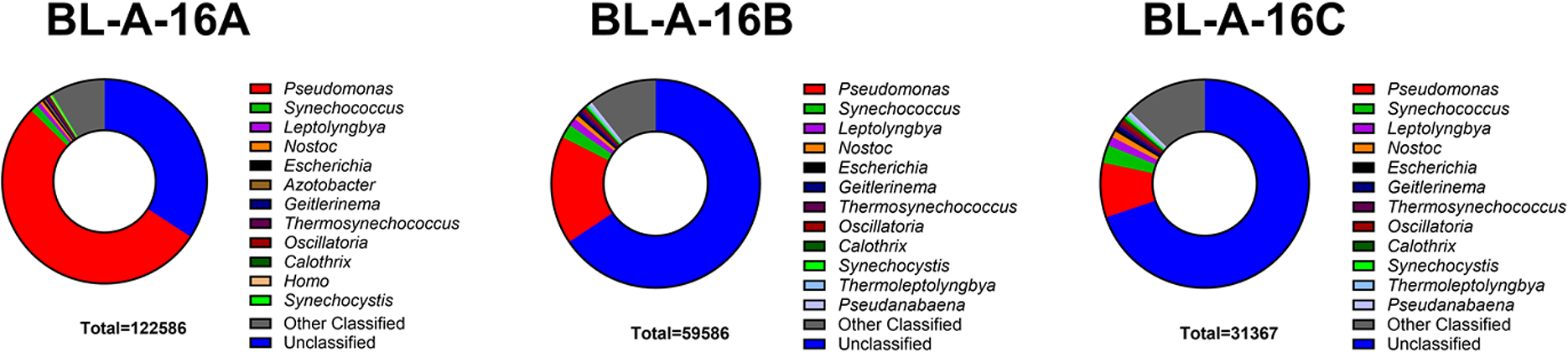
The EPI2ME classification of reads from sequencing runs of BL-A-16. The EPI2ME platform seemed to struggle with classifying cyanobacterial DNA, notice the large fraction of unclassified reads and numerous different cyanobacterial genera annotated for this single species among classified reads. However, the decrease in *Pseudomonas* reads during the iterative lysis step is clear, dropping from 53% to 17% to 8% of total reads from fraction A to B to C shows clearly the winnowing of heterotrophic DNA from the sample during the iterative lysis.

### Genome assembly using the combined genomic data

We used the Porechopper and FiltLong tools to trim and filter our basecalled nanopore reads before assembly with Flye to afford metagenomic assemblies of our cyanobacterium/heterotrophic microbiome communities. In both the assemblies of BL-A-14 and BL-A-16, the largest of the contigs was a closed circular chromosome which based on BLAST of representative genes (i.e. the 16S gene) we suspected was our cyanobacterial host (SI Tables A and B). For BL-A-14 seven circular plasmids were also assembled, while the BL-A-16 assembly had two circular plasmids. Because of the higher error rate of nanopore sequencing versus other sequencing platforms, we first used Medaka correction with the original nanopore reads and then Polypolish with the Illumina reads to polish this genome.[30,31] DFAST annotation of these cyanobacterial chromosomes showed that the coding ratio increased after polishing while the total number of coding sequences (CDS) decreased (SI Table C); due to what we believe is correction of errors, such as homopolymer errors, within genes that incorrectly introduced stop codons within a single reading frame leading to two split CDS for a single gene in the early drafts of the assembly. A Type Strain Genome Server (TYGS) comparison of the BL-A-14 chromosome suggested this strain was a new species and showed a clade with our strain as well as *Stenomitos rutilans* HA7619-LM2 and *Neosynechococcus sphagnicola* CAUP A 1101 by 16S compared to a clade with more distantly related organisms such as *Ensifer adhaerens* ATCC 33212 or *Agrobacterium burrii* RnrT by whole genome comparison (SI Figures D and E), a confusing result that made little sense based on how divergent these near neighbors were. By using a simpler BLAST comparison of the 16S gene from our BL-A-14 chromosome we saw the top hit was to *Leptolyngbya* sp. CENA387 (KR137608) with a 99.6% pairwise identity; comfortably assigning our strain as a *Leptolyngbya* sp. While there are complete chromosome assemblies for the *Leptolyngbya* genus, they vary in size from 4.4 to 7.2 Mb (see ASM2624063v1 and ASM236825v1), so we hypothesize that the discrepancy between genome-based and 16S-based analysis is the difficulty of genomic comparison with large size differences in the genome. Similarly, the TYGS assigned the BL-A-16 chromosome as a new species (SI Figures F and G), while a BLAST comparison showed a top hit to *Limnothrix* sp. SK1-2-1 (LC272581) with a 99.9% pairwise identity between their 16S genes. In this case we found only ten genomes of this genus on NCBI’s genome database, none of which were assembled to the chromosome level, explaining why the TYGS’s genome-to-genome comparisons failed to recognize similarity to species in this genus.

## Conclusions

Our lab’s interest in genomic investigation of cyanobacteria collected from Nature faces some of the hurdles long known in this field, such as the challenge of having heterotrophic DNA contaminate DNA extractions from a cyanobacterial strain serving as host to heterotrophic bacteria. [14,18–20] Our initial attempts to use Illumina sequencing for our strains was confounded by just this result, leading to incomplete and disjointed assemblies of the cyanobacterial genome. Utilizing a modified lysis approach where we used iterative steps beginning with a quick lysis ill-suited to lyse the cyanobacterial cells, followed by longer lysis steps repeated on the residual cell pellet we saw heterotrophic reads of the sequenced DNA overrepresented in DNA from the first lysis step while cyanobacterial reads were enriched in later fractions. To derive DNA suitable for nanopore sequencing we had to rely on multiple bead-based size selection steps to enrich for longer molecules, which did allow us to utilize this sequencing technology. However, we believe that the ability to target a specific portion of the microbiome could be broadly applicable to many different sequencing technologies. In the remarkable change in proportion of total reads coming from *Psuedomonas* in the DNA extracts of *Limnothrix* sp. BL-A-16, from 54% in fraction A to 9% in fraction C (as assessed by EPI2ME analysis) we see clear evidence that our strategy can help pull apart complex microbiomes during the DNA extraction step. Other researchers interested in cyanobacterial genomics could similarly apply this workflow to their own projects. By purposely sequencing the fraction A we may also be able to investigate the composition of the microbiome which is challenging where culturing methods can prove insufficient to isolate each member of the microbiome.[18] Where other methods of isolating a cyanobacteria-enriched DNA fraction are destructive to the DNA of the heterotrophic microbiome, our new methods recovers this DNA.[14] We are currently working to use our method to whole genome sequence other strains in our lab’s culture collection.

## Materials and Methods

### General Experimental Equipment and Materials

The autoclave used for sterilization was a Steris™ AMSCO^®^ C Series Remanufactured Small Steam Sterilizer. Long-term storage of samples was done at ultra-low temperatures in a VWR® Ultra-Low Temperature Upright Freezer set to -70 °C. The LED grow lights from Gardener’s Supply Company™, with height adjusted for cultures to receive 1550 lux on a 16:8 light/dark cycle timer. A ZEISS Widefield Axio microscope from the Glycoscience Center of Research Excellence (GlyCORE) Imaging Research Core was used to image the cyanobacterial and algal cultures. The lyophilizer used was a Labconco^®^ Freeze Dry System/Freezone 2.5. The 50 mL tubes were centrifuged in a Sorvall™ Legend™ XTR Fixed Angle Centrifuge. The microcentrifuge used was a Sorvall™ Legend™ Micro 21R Microcentrifuge. The heat block used was a Fisherbrand™ Mini Dry Bath. The vortex used was a Fisherbrand™ Science Industries Inc., Vortex Genie 2. An Invitrogen™ Qubit™ 4 Fluorometer was used in accordance with the manufacturer’s Quick Reference Qubit™ Assays Qubit protocol. A Thermo Scientific™ mySPIN™ 6 Mini Centrifuge was used to spin samples in the 1.5 mL microcentrifuge and PCR tubes down. The nanodrop was a Thermo Scientific NanoDrop One^C^.

The petri dishes used were 15x100 mm Fisherbrand™ Petri Dishes with Clear Lid. The 50 mL and 15 mL polypropylene conical natural centrifuge tubes were Falcon™ brand. The inoculation loops used were Fisherbrand™ Disposable Inoculating Loops and Needles, Flexible Needle and Loop. The 1.5 mL microcentrifuge tubes used were VWR^®^

Polypropylene Microcentrifuge Tube. For column binding, the Zymo Research® DNA Clean and Concentrator™-5 kit according to the protocol from the manufacturer for genomic DNA (>kb) was used. The isopropanol (IPA) used was LabChem™ Isopropyl Alcohol, LCMS grade. The ethanol used was Decon Laboratories™ 200 proof ethanol, anhydrous that meets USP specifications. Beckman Coulter SPRI™ size selection beads were used in accordance with the Left Side Size Selection SPRIselect protocol according to the manufacture’s guidelines. Chemicals were purchased from high quality lab suppliers, specifically, the germanium (IV) dioxide was ≥99.99% trace metals basis from Aldrich, the nystatin was from Alfa Aesar.

### Isolation of Cyanobacteria, Collection of Cyanobacterial Cells

FW-BG-11 plates were made according to a recipe adapted from the University of Texas at Austin Culture Collection of Algae’s BG-11 media recipe.[29] When making solid agar plates, after autoclave sterilization, just before pouring the cooled medium, 200 μL of a 20 mg/mL stock solution of germanium dioxide in water, filter sterilized, and 1 mL of a 50 mg/mL stock solution of nystatin in ethanol was added to the medium.

Cyanobacterial strains were isolated from fresh water sources in Northern Mississippi. BL- A-14 was collected from Enid Lake (assigned BioSample number SAMN42824079); BL-A- 16 was collected from the Tennessee-Tombigbee Waterway (assigned BioSample number SAMN42824080). Water samples were collected in sterile 50 mL centrifuge tubes, placed on ice, and then brought to the lab. 50 μL of the water samples were plated directly as well as a 10x concentrated sample prepared by centrifugation of 1000 μL of the water sample at 21.0 g at 13 °C for 5.0 min then removing the top 90% of the water sample before vortexing and plating 50 μL of the concentrated material.

All plates were placed under the grow lights and allowed to grow for 1-2 months depending on when the cyanobacterial colonies appear and were large enough to be picked. Once colonies appeared, colonies with different morphologies were picked from the plate and re-struck to a fresh plate under sterile conditions. This process was repeated until a single cyanobacterial morphology was observed on the plate. Then from that single morphology plate a colony was picked into 125 mL autoclaved Erlenmeyer flasks with 50 mL of liquid FW-BG-11 medium. Cultures were then grown in continuous culture for circa two months between passages with 1 μL/mL inoculation into 50 mL of fresh media.

Cultures were harvested when the culture has substantial biomass, before bleaching or yellowing was observed. Once the culture was ready for harvesting, it was poured in to a 50 mL centrifuge tube. The culture was then centrifuged for 15 minutes at 10,000 g at 13 °C. After centrifugation the biomass accumulated in the pellet was frozen at ultra-low temperature. The frozen pellet was then lyophilized overnight. Once the pellet was fully dried, it was either placed back into the ultra-low temperature freezer for storage until use, or used immediately.

### DNA Isolation for Illumina Sequencing

For BL-A-14, after collecting cyanobacterial pellets from 50 mL cultures, gDNA isolation was performed using the OMEGA Bio-Tek E.Z.N.A.^®^ Plant DNA kit according to the manufacturer’s instructions. In brief, the lyophilized cyanobacterial cells underwent lysis employing P1 buffer, while the P2 buffer was utilized to precipitate proteins and polysaccharides. Subsequently, the resulting cleared supernatants were then bound to a HiBind column, washed with the manufacturer’s buffers, and eluted using an elution buffer. For BL-A-16, the DNA was isolated using OMEGA Bio-Tek E.Z.N.A.^®^ Bacterial DNA kit following the manufacturer’s protocol using the optional bead beating by vortexing with ca. 25 mg of glass beads for 5 min. DNA quantification was carried out via nanodrop before shipping for sequencing, where DNA concentration was insufficient for the vendor (in the case of BL-A-16), the material was concentrated using Zymo’s DNA Clean & Concentrator-5 kit according to the manufacturer’s protocol. This material was sent for Illumina MiSeq 2x150 bp sequencing with a commercial vendor (Genewiz).

### Iterative Lysis Protocol for DNA Isolation Targeting Cyanobacterial DNA

Two lyophilized pellets from two 50 mL harvests of cultures of the desired strain were used for the DNA isolation. An adapted version of the OMEGA Bio-Tek E.Z.N.A.® Plant DNA Kit protocol was followed: First, the two dried cell pellets were combined and 1000 μL of P1 buffer was added to the pellets, this mixture was vortexed briefly. Once vortexed, the 50 mL centrifuge tube was placed in a 60-65 °C water bath for 10 minutes. After the 10-minute incubation, the lysate was transferred to a 1.5 mL microcentrifuge tube and centrifuged at 21.0 g at 13 °C for 2.0 minutes in the microcentrifuge. Once centrifuged, the supernatant was collected by pipette and placed in a separate sterile 1.5 mL microcentrifuge tube.

With the pellet from the previous step, 800 μL of P1 buffer, 10 μL of lysozyme, 10 μL of Proteinase K and ca. 10-20 mg of glass beads were added to the sample; we vortexed this mixture briefly. The sample was then placed in the preheated heat block at 65 °C for 20 minutes, we inverted the tube six times halfway through the incubation to mix. Then the supernatant was collected as before. The second pellet was incubated with 800 μL of P1 buffer, 10 μL of lysozyme, and 10 μL of Proteinase K for one hour at 65 °C and inverted six times twice, at the 20- and 40-minute points. After the incubation, the sample was centrifuged as before, and the final supernatant was collected.

With the supernatants from the previous steps, 140 μL of P2 buffer was added and inverted six times to mix. The mixture was then centrifuged at 21.0 g at 13 °C for 10 minutes in the microcentrifuge. Once centrifuged, the supernatant was collected in a sterile 1.5 mL microcentrifuge tube and 70% the volume of IPA was added. The tube was then inverted six times to mix before centrifuging for 5 minutes at 21.0 g and 13 °C in the microcentrifuge. Once centrifuged, the supernatant was discarded, collecting the pellet.

Next, 300 μL of 65 °C MiliQ water was added to the pellet, to aid the pellet dissolving, it was placed in the pre-heated 65 °C heat block for 10 minutes. After the incubation, 4.0 μL of RNase was added and the DNA was measured on the Qubit with 2.00 μL of the sample.

Next, the samples were cleaned and concentrated during the column binding step using the DNA Clean and Concentrator kit in accordance with the protocol for genomic DNA from the manufacturer. For the elution steps, we specifically used first 17.0 μL of sterilized MiliQ added directly to the column matrix and then incubated at room temperature for 5 minutes before collecting this eluent by centrifugation for 30 seconds at 21.0 g and 13 °C. The second elution was then performed with 15.0 μL of sterilized MiliQ water and a 3-minute incubation before being combined with the first again by centrifugation. The column was then discarded, and the sample was then measured on the Qubit as above. Once measured, the sample was frozen at -70 °C for storage.

### Bead-Base Size Selection of DNA Before Nanopore Sequencing

Following the SERAMAG left side size selection protocol from the manufacturer, we added beads at a 2:1 ratio to the sample volume. Next the 1.5 mL microcentrifuge tube was flicked for 60 seconds to mix and avoid shearing DNA before being briefly spun down and incubated at room temperature for 7 minutes. The tube was then placed on the magnetic rack for 5 minutes to pellet the beads fully. Then the supernatant was pipetted off and discarded. The beads were washed twice, leaving the tube on the magnet, with 200 μL of freshly prepared 85% ethanol, taking care not to disturb the pellet. After the second wash, the sample was carefully aspirated and as much of the wash solution as possible was removed before drying the beads on the magnet for 7 minutes at room temperature. The microcentrifuge tube was removed from the magnetic rack and 50 μL of TE (Tris-EDTA) buffer was added to elute the DNA. The SERAMAG beads were resuspended by flicking the tube for 60 seconds, the tube was then briefly spun down before being incubated for 7 minutes at room temperature. After incubation, the microcentrifuge tube was placed in the magnetic rack for 5 minutes to fully settle the beads. Once settled, the supernatant was carefully pipetted off and retained in a new sterile microcentrifuge tube.

The entire SERAMAG size selection protocol was repeated a second time on this bead-purified material and then the final recovered DNA was measured on the Qubit as before. The sample was then frozen at -70 °C until use.

### Genome Sequencing via the Nanopore Platform

Doubly size selected DNA was sequenced on an Oxford Nanopore Flongle cells (FLG001) using the Ligation Sequencing Kit (SQK- LSK100) according to the manufacturer’s protocol. The end-prep and nick repair were conducted without deviation from the protocol using 500 ng of the doubly size selected DNA. Next during the adapter ligation, again the manufacturer’s protocol was followed using New England Biolab’s Quick T4 DNA Ligase, however, during the cleanup step instead of washing the AMPure XP beads with 125 μL of Long Fragment Buffer or Short Fragment Buffer, we first washed with 125 μL of Long Fragment Buffer then 125 μL of Short Fragment Buffer for the second wash. During the final loading of the Flongle, we made sure to use reagents in glass vials from the Flongle Sequencing Expansion, following the manufacturer’s protocol. In setting the run parameters we selected our flow cell, kit, and a 48-hour total run time.

### Bioinformatic Processing and Analysis of Nanopore Reads

Illumina data was processed by the commercial vendor; briefly, after a FastQC quality check *de novo* genome assembly was carried out with Spades v 3.10 and genome statistics were generated via QUAST.

After the nanopore sequencing run, data was transferred to the computing resources in the GlyCORE Computational Chemistry and Bioinformatics Core for processing and assembly of the genomes. The first step of processing the sequencing reads was to basecall the .fast5 nanopore reads using Guppy basecaller (version 6.4.2), for which we used the super accuracy configuration file for our kit and flowcell. Once the reads were basecalled to .fastq files, a pycoQC report was performed to assess the quality of reads [32].

The EPI2ME analysis was conducted with the software available from Oxford Nanopore [33]. We submitted our basecalled .fastq files to “Fastq WIMP” (What’s In My Pot) analysis. We set the parameters to Min q score of 7, with no detect barcode function, a minimum length filter of 500 bp, and a max length filter of zero.

We used the Porechop tool to remove adapter sequences from the raw basecalled reads [34]. Next, these adapter-free sequences were run through the FiltLong tool [35]. In the FiltLong settings, we put the length cutoff threshold to the tenth percentile as determined by the pycoQC report or 500 bp, whichever was shorter. We set the quality score to filter out at least 1% of reads by quality, but as this step occurs after filtering by length, reads were effectively only filtered by length.

### Genome Assembly, Statistics, and Comparison

Once Filtlong was complete, an initial draft assembly was constructed with Flye using the metagenomic setting (version 2.9) [36].

Medaka was used to improve the draft Flye assembly [37]. For the Medaka run the inputs were the draft Flye assembly and the combined reads files after FiltLong trimming, the default batch size was 500 using model r941_min_sup_g507. After Medaka, the Polypolish tool was run to correct any of the long-read assembly errors with previously collected Illumina data [30,31]. Polypolish was coupled with the BWA, Burrows-Wheeler Alignment tool [38]. BWA was used to index the Medaka-corrected consensus assembly to the Illumina reads as a .sam alignment. Polypolish was then run using the Medaka consensus assembly file, and the two filtered .sam files to produce a polished assembly of our sequenced genomes.

DFAST™ annotation was done with the online version of the tool available from the DNA Data Bank of Japan, National Institute of Genetics [39,40]. In this project, we ran the DFAST tool on our circular chromosomes of cyanobacterial DNA extracted from our metagenomic assemblies.

We used the TYGS to analyze our cyanobacterial chromosomes and compare them to the TYGS’s database, based on both 16S and whole genome-based phylograms [41]. Each polished chromosome was uploaded to the server as a separate job with the phylogram results presented in the SI (see SI Figures D-G).

Finally, the other contigs present within the assembly were analyzed. Two contigs across the assemblies BL-A-14-contig_10 and BL-A-16-contig_7 were nearly identical to each other and had a >99% pairwise identity with *E. coli* strain Q4552 plasmid pECQ4552_IHU08 (CP077071) as assessed using a blastn search in Geneious prime (version 2023.0.4). Given this similarity across to the two assemblies, these contigs were deemed contaminants from our DNA isolation workflow; note that the coverage of these contigs was also far higher than any other contig in the assemblies (see SI Table A and B). In the assembly of BL-A-14, 34 contigs between 1,216 and 6,561 bp were binned out by their high GC content (ranging from 68.2% - 76.2%). Again, blastn analysis showed the nearest hits to these nucleotide sequences were not cyanobacterial sequences, instead these hits were all Actinobacteria, suggestive that these contigs came from a heterotrophic member of *Leptolyngba* sp. BL-A-14’s microbiome. The remaining seven contigs were assembled as circular contigs between 2,182 and 288,243 bp, with GC content far closer to the cyanobacterial genome (ranging from 48.3% to 52.0%), and blastn analysis of open reading frames on these contigs showed the most similar sequences were mostly cyanobacterial. As such we assigned these seven contigs as plasmids of this strain. We were not surprised, based on our EPI2ME analysis, in finding several contigs within the assembly of BL-A-16 that were more similar to *Pseudomonas*. Binning by GC content pulled five contigs out of the assembly, with lengths of 182,682 to 1,612,854 bp and GC content ranging from 62.4% to 63.6%. Blastn analysis of open reading frames within these contigs confirmed them as being *Pseudomonas*-derived, we even found three identical copies of the 16S gene which had a 100% pairwise identity to the full 16S gene of *Pseudomonas argentinensis* strain AFS0086324 (OP986251). Two of the remaining contigs were assembled as circular by Flye, BL-A-16-contig_10 had a length of 55,181 bp and a GC content of 55.3% while BL-A-16-contig_11 had a length of 15,919 bp and GC content of 53.0%. We assigned these sequences as cyanobacterial plasmids based on their GC content and circular topology. Blastn analysis of open reading frames from within BL-A-16- contig_11 confirmed this assignment with hits to other cyanobacterial sequences, while BL- A-16-contig_10 showed few hits to any known sequences in GenBank (cyanobacterial or otherwise). The last contig was a linear 5,804 bp 51.3% GC fragment which had no blastn hits to known sequences so this fragment was removed from the assembly.

The polished assemblies of these two species were deposited with GenBank under the BioProject number PRJNA1048477. The assembly of *Leptolyngbya* sp. BL-A-14 was given the accession numbers CP166616-16623 (chromosome as CP166621), while the assembly of *Limnothrix* sp. BL-A-16 was given the accession numbers CP166613-16615 (chromosome as CP166615). The full PycoQC reports of the nanopore runs, the scaffolds of the vendor’s Illumina-based assemblies, and the original Polypolished assemblies (before removal of contigs assessed as being from the heterotrophic microbiome by GC content and BLAST similarity) are available on the eGrove archive at https://egrove.olemiss.edu/pharmacy_facpubs/290/.

## Acknowledgements

We thank both the Imaging Research Core and the Computational Chemistry and Bioinformatics Research Core within the University of Mississippi’s Glycoscience Center of Research Excellence (NIH Project Number 5P20GM130460) for use of their equipment and facilities as outlined in the experimental sections above. We thank the Stevens lab in our department for access to and use of their Nanodrop. We thank the Center of Biomedical Research Excellence in Natural Products Neuroscience (COBRE-NPN) for use of their MiliQ system. Funding to support this project came from a startup package provided by the University of Mississippi Provost’s office and the Department of BioMolecular Sciences in the School of Pharmacy to Dr. Boudreau; as well as a Pilot Project subaward to Dr. Boudreau’s lab from the COBRE-NPN, the COBRE-NPN award is funded by the National Institute of General Medical Sciences (NIGMS) at the National Institutes of Health as one of its Centers of Biomedical Research Excellence (Grant Number P30GM122733). The content is solely the responsibility of the authors and does not necessarily represent the official views of the National Institutes of Health.

